# L-Dopa incorporation into tubulin alters microtubule dynamics and reduces dendritic spine invasion and synapse maintenance

**DOI:** 10.1101/2024.05.20.595021

**Authors:** Agustina Zorgniotti, Aditi Sharma, Sacnicte Ramirez-Rios, Chadni Sanyal, Martina Aleman, Yanina Ditamo, Marie-Jo Moutin, C. Gastón Bisig, Leticia Peris

## Abstract

Previous studies have shown that L-Dopa, a tyrosine analog used in Parkinson’s disease treatment, can be incorporated into α-tubulin C-terminal tail via the tubulin tyrosine ligase (TTL) and polymerize into microtubules. In this work, we demonstrated that mature wild type hippocampal neurons treated with L-Dopa exhibited reduced dendritic spine density, primarily affecting mature dendritic spines. In these neurons, L-Dopa treatment significantly reduced tyrosinated α-tubulin levels without altering detyrosinated or Δ2 α-tubulin levels, suggesting the formation of a new tubulin pool, likely composed of L-Dopa-α-tubulin.

In vitro analysis of the activity of the purified VASH1-SVBP complex, the most abundant tubulin carboxypeptidase in brain, revealed that L-Dopa incorporation into α-tubulin modified the binding of the complex to microtubules and reduced its carboxypeptidase activity. These results suggest that L-Dopa incorporation into tubulin alters the properties of microtubules and affects their ability to interact with the enzyme.

To confirm the implication of L-Dopa-microtubules in dendritic spine alterations observed in wild type neurons, we analyzed the effect of L-Dopa treatment in neurons lacking the enzymes of the α-tubulin detyrosination/tyrosination cycle. In these cells, L-Dopa cannot be incorporated into α-tubulin due to the absence of the ligase (in TTL KO neurons) or the reduction of detyrosinated α-tubulin levels (in SVBP KO neurons). L-Dopa treatment did not modify dendritic spine density in TLL KO or SVBP KO neurons, clearly demonstrating that the alterations in dendritic spines seen in WT neurons are due to the incorporation of L-Dopa into tubulin.

Further analysis revealed that L-Dopa treatment decreased the percentage of spines containing excitatory synapses in wild type neurons, but not in TTL KO or SVBP KO neurons, suggesting a cumulative synaptic defect due to L-Dopa incorporation into microtubules. Additionally, L-Dopa altered microtubule dynamics by increasing catastrophe frequency and reducing comet lifetime, which led to fewer microtubules entering dendritic spines and decreased spine resistance to pruning.

Taken together, our results demonstrate that L-Dopa incorporation into α-tubulin drastically affects synaptic homeostasis, reaffirming the importance of balanced detyrosination/tyrosination of tubulin within the synaptic compartment. The abnormal dynamics of L-Dopa-microtubules and the reduction of dendritic spines and excitatory synapses highlight a novel mechanism of L-Dopa-induced synaptotoxicity.

## INTRODUCTION

Parkinson’s disease (PD) is a neurodegenerative disorder characterized by progressive degeneration of dopaminergic neurons located in the midbrain, leading to slowing of movement, rigidity, and tremor, in addition to cognitive problems an increased risk for dementia and depression (Poewe et al., 2017; Woods & Tröster, 2003). Administration of L-3,4-dihydroxyphenylalanine (L-Dopa) is the gold-standard pharmacological therapy. L-Dopa, the immediate precursor of dopamine, mitigates motor symptoms; however, its use is limited by the development of abnormal dyskinetic movements (L-Dopa-induced-dyskinesia; LID) and affective and cognitive disorders after long-term treatment (Bastide et al., 2015; Marano et al., 2019). The pathophysiology of motor complication associated with Parkinson’s disease, particularly L-Dopa-induced-dyskinesia, has been extensively investigated (Bastide et al., 2015). However, despite several molecular-level studies of this phenomenon, the cause of L-Dopa-induced-dyskinesia remains unknown.

Dendritic spines are considered the smallest anatomical structure where the biochemical and electrophysiological signals of the glutamatergic and dopaminergic synapses are integrated (Chen & Sabatini, 2012). Increasing evidence suggests that spines dynamically change their density and morphology, and that these changes are deeply involved in synaptic plasticity (Kasai et al., 2010). In various experimental models, as well as in studies in Parkinson’s disease patients, it has been observed that prolonged treatment with L-Dopa generates alterations in homeostatic synaptic mechanisms. These alterations have been related with changes in the synaptic density in medium spiny neurons, in their functionality and structure, and may form the basis of L-Dopa-induced complications in patients with Parkinson’s disease (Nishijima et al., 2018; Picconi et al., 2018). The emerging consensus is that alteration of dopamine release and reuptake and abnormalities in neurotransmitter systems are involved in L-Dopa-induced-dyskinesia development (Fabbrini & Guerra, 2021). Regarding structural changes, rat models of L-Dopa-induced-dyskinesia have shown a critical association between L-Dopa administration and abnormal synaptic plasticity at corticostriatal synapses (Picconi et al., 2003), as well as a decrease and enlargement of dendritic spines in striatal medium spiny neurons (Nishijima et al., 2013).

In neurons, the microtubule (MT) cytoskeleton composed of α- and β-tubulin, plays a key role in the development and maintenance of axons and dendrites. The microtubule dynamics, critically contribute to synaptic structure and function within both pre- and postsynaptic compartments (Waites et al., 2021). Dynamic microtubules differ from stable microtubules in their capacity for stochastic transitions, shifting between depolymerization and polymerization, and vice versa, a process named dynamic instability (Avila, 1990; Mitchison & Kirschner, 1984)). These dynamic microtubules regulate synaptic vesicle cycling by generating pathways for bidirectional transport between presynaptic terminals, a rate-limiting step in exocytosis at the release sites (Guillaud et al., 2017; Piriya Ananda Babu et al., 2020; Qu et al., 2019). In dendritic spines, while the cytoskeletal structural core is formed by actin filaments, dynamic microtubules originating from the dendritic shaft sporadically invade into the spine head and directly influence the regulation of spine composition and morphology. Microtubule entry into spines is dependent on synaptic activity, Ca^2+^ influx, actin polymerization, post-translational modifications of tubulin (PTMs) and correlates with changes in synaptic strength (Dent, 2020; McVicker et al., 2016; Peris, Parato, Qu, Soleilhac, Lante, et al., 2022). Tubulin PTMs preferentially accumulate on stable microtubules. The combinatorial nature of these covalent modifications, as well as many different tubulin isotypes, give rise to a ‘tubulin code’ that regulates a variety of neuronal functions (Janke & Magiera, 2020). The wide variety of post-translational modifications of tubulin controls the interaction with motor proteins, microtubule severing enzymes, microtubule-associated proteins (MAPs) and microtubule end-binding proteins (Balabanian et al., 2017; Janke & Kneussel, 2010; Peris et al., 2006, 2009). These events, in turn, regulate the flux of vesicles, organelles, RNA and multiprotein complexes into and out of synapses (Guedes-Dias & Holzbaur, 2019).

Our series of studies in the 1970s demonstrated that the α-tubulin C-terminus is subject to post-translational cyclic detyrosination/tyrosination (Arce et al., 1978; Barra et al., 1973, 1974, 1988). Upon microtubules assembly, the tyrosine residue encoded at the C-terminal α-tubulin (Tyr-tubulin) is removed by the carboxypeptidase enzymes to generate detyrosinated tubulin (deTyr-tubulin) (Arce & Barra, 1983). These enzymes include the heterodimeric complexes composed of vasohibins (VASH) 1 and 2, along with the small vasohibin-binding protein (SVBP) (Aillaud et al., 2017; Nieuwenhuis et al., 2017), and the recently described microtubule associated tyrosine carboxypeptidase (MATCAP) (Landskron et al., 2022). Following release of tubulin from microtubules, tubulin tyrosine ligase (TTL) rapidly tyrosinates deTyr-tubulins and, after their re-assembly into tyrosinated newly formed microtubules, the cycle continues (Ersfeld et al., 1993). The pool of deTyr-tubulin can be further processed by cytosolic carboxypeptidases that remove the terminal glutamate, generating mainly Δ2-tubulin (Paturle-Lafanechère et al., 1991). Unlike deTyr-tubulin, Δ2-tubulin cannot be re-tyrosinated by the TTL enzyme (Paturle-Lafanechère et al., 1994; Prota et al., 2013; Rüdiger et al., 1994).

Our previous studies have shown that L-Dopa can be incorporated into the α-tubulin C-terminus in vitro and in living cells (Dentesano et al., 2018; Rodriguez et al., 1975; Zorgniotti et al., 2021), as well as other tyrosine analogues (Arce et al., 1978; Barra et al., 1973; Bisig et al., 2002; Ditamo et al., 2016; Purro et al., 2003). Most of the analogues studied to date can be cyclically incorporated and released from ɑ-tubulin, completing the detyrosination/tyrosination cycle, which is not the case of L-Dopa. Indeed, we demonstrated that once L-Dopa is incorporated into the α-tubulin C-terminus, it cannot be released by tubulin carboxypeptidases under conditions that allow rapid release of tyrosine from Tyr-tubulin (Zorgniotti et al., 2021). Treatment of neuron-like cells with L-Dopa and consequent increase in L-Dopa-tubulin resulted in altered microtubule dynamics in neurite-like processes (Dentesano et al., 2018). In cultured hippocampal neurons treated with L-Dopa, we observed that mitochondrial traffic was altered due to a reduced capacity of KIF5B to bind to L-Dopa-microtubules (Zorgniotti et al., 2021).

Tyr-tubulin is associated with dynamic microtubules as tyrosinated microtubules are sensitive to the microtubule-depolymerizing drug nocodazole (Ahmad et al., 1993; Bré et al., 1987; Kreis, 1987) and are substrates for Kinesin-13 depolymerizing motors (Peris et al., 2009). In contrast, deTyr- and Δ2-tubulins are associated with more stable, longer-lived microtubules (Kreis, 1987; Webster et al., 1987). The role of tubulin detyrosination/tyrosination in many cell physiological processes remains poorly understood; however, there is increasing evidence of its importance in various specialized microtubule functions (Pagnamenta et al., 2019). Presence or absence of a C-terminal tyrosine residue on α-tubulin affects processivity of kinesins and dyneins motors (Gundersen et al., 1984; Gundersen & Bulinski, 1986; Verhey & Gaertig, 2007), the binding of MAPs (Bieling et al., 2008; Peris et al., 2006), and chromosome movements during mitosis (Barisic & Maiato, 2015). In neurons, the relative proportions of Tyr- and deTyr-tubulin, display temporal and spatial variations during neuronal differentiation, and these changes correlate with alterations in microtubule dynamics. In mature cultured neurons, microtubules become more stable and accumulate PTMs. Tyrosinated microtubules are found in the outer part of dendrites, whereas stable acetylated and Δ2-tubulin bundles are located in the inner part (Tas et al., 2017). Dynamic tyrosinated microtubules from the outer part of the dendrites transiently invade dendritic spines in an activity-dependent manner, suggesting that microtubules, in cross-talk with the actin cytoskeleton, play an important role in synapse function and plasticity (Dent et al., 2011; Hu et al., 2008; Jaworski et al., 2009; Schätzle et al., 2018).

A balanced detyrosination/tyrosination tubulin cycle is essential for neurons and brain integrity. In TTL knockout mice, the complete loss of the tyrosination enzyme results in lethal brain and neuronal defects (Erck et al., 2005; Hosseini et al., 2022; Marcos et al., 2009; Martínez-Hernández et al., 2022; Peris et al., 2006, 2009; Sanyal et al., 2023). Partial reduction of the enzyme in TTL heterozygous mice exhibit decreased dendritic spine density and both synaptic plasticity and memory deficits (Peris, Parato, Qu, Soleilhac, Lanté, et al., 2022). On the other hand, the absence of VASH-SVBP carboxypeptidases, which leads to defective tubulin detyrosination, causes structural brain abnormalities and cognitive deficiencies in both humans and mice (Iqbal et al., 2019; Pagnamenta et al., 2019). The synaptic function and disruption in this balance has been recently shown associated with neurodegeneration in Alzheimer’s disease (Peris, Parato, Qu, Soleilhac, Lante, et al., 2022).

Surprisingly few studies exist on how the regulation of microtubule dynamics through tubulin modifications affects the neurotransmitter machinery of synapses and the resulting impact on neuronal degeneration. Here we show that L-Dopa incorporation into microtubules leads to a reduction in dendritic spine density in wild-type hippocampal neurons, without altering spine morphology. Interestingly, this was not observed in neuronal cultures of knockout models lacking detyrosination/tyrosination cycle enzymes, indicating that the defects observed in wild type neurons are exclusively due to the presence of L-Dopa in microtubules. We found that the presence of L-Dopa on tubulin disrupts the binding to microtubules and the activity of the most abundant carboxypeptidase in the brain, VASH1-SVBP. We also demonstrate that L-Dopa incorporation into microtubules diminishes excitatory synapses and modifies global microtubule dynamics, thereby reducing microtubule invasion into dendritic spines, which makes them more susceptible to spine pruning.

These findings are relevant to Parkinson’s disease patients under long-term L-Dopa treatment. Our results suggest that, concomitantly with beneficial enhancement of dopamine synthesis, the tyrosination state of tubulin (and microtubules) is gradually altered by incorporation of L-Dopa into α-tubulin C-terminus. This could affect key processes for the maintenance of synaptic connectivity and functionality and be responsible for some of the neurological defects associated with prolonged L-Dopa therapy.

## MATERIAL AND METHODS

### Animals

All experiments involving mice were conducted in accordance with the policy of the Institut des Neurosciences de Grenoble (GIN) and in compliance with the French legislation and European Union Directive of 22 September 2010 (2010/63/UE). TTL heterozygous (TTL+/-) mice and mice homozygous for an inactivated small vasohibin binding protein allele (referred to as SVBP-KO), were obtained as previously described (Erck et al., 2005; Pagnamenta et al., 2019) and maintained in a C57BL/6 genetic background by recurrent back-crosses with C57BL6 animals from Charles River Laboratories.

### Genotyping

PCR amplifications were performed on alkaline lysates of toe clips or tail cuts of E18.5 mouse embryos. Briefly, mouse tissue was incubated for 30 min at 95°C in alkaline solution (NaOH 25 mM, EDTA 0.2 mM, pH 12.0). Neutralization was performed by adding 40 mM Tris, pH 5.0. Lysates were then analyzed by PCR with corresponding primers and Econo Taq PLUS Green Mix (Euromedex). Primers pairs for testing the TTL mouse strain were 5 ′ -GGCGACTCCATGGAGTGGTGG-3 ′ and 5 ′CCCAACAT-CACATTCTCCAAATATCAAAG-3 ′ (TTL wild type, 1032 bp) and 5 ′GATTC-CCACTTTGTGGTTCTAAGTACTG 3′and 5′CCCAACATCACATTCTCCAAATATCAAAG-3 ′ (TTL KO, 900 bp). The four primers were used in a single reaction (Fig. S1A). Primers pairs for testing SVBP mouse strain were 5 ′GATCCACCTGCCCGGAAA 3 ′ and 5 ′TTTCTTCCAGCACCCTCTCC 3 ′ (SVBP wild type, 170 bp) and 5 ′TTTCTTCCAGCACCCTCTCC 3 ′ and 5 ′CAAACCATGGATCCACGAAA 3 ′ (SVBP KO, 167 bp). These latter reactions were done separately as in Konietzny 2024. The following amplification protocols were used: TTL, 95°C for 5 min, 35 cycles of [95°C for 1 min, 50°C for 1 min, 72°C for 1 min], 72°C for 2 min; SVBP, 95°C for 5 min, 33 cycles of [95°C for 30 s, 50°C for 30 s, 72°C for 30 s], 72°C for 2 min. DNA was analyzed on 1.2% and 2% agarose gels for TTL and SVBP, respectively.

### Plasmids

For lentiviral experiments, a vector eGFP-pWPT (Addgene #12255, kind gift from D. Trono) was used to express eGFP and a home-made vector containing LifeAct-RFP – IRES - EB3-YFP were used to express both LifeAct-RFP and EB3-YFP. For the last one, the cDNAs encoding LifeAct (aa 1-17 of Saccharomyces cerevisiae ABP140, NP_014882) C-terminally fused to RFP, and encoding human EB3 (NP_001289979) C-terminally fused to YFP were cloned in pLV-mCherry vector During the cloning process, mCherry cDNA was removed and replaced by an IRES (encephalomyocarditis virus internal ribosome entry site) sequence to allow EB3-YFP expression. PCR amplification and cloning of cDNAs was performed with Phusion DNA polymerase (Thermo Scientific) and In-Fusion HD Cloning kit (Clontech), respectively. The construct was verified by sequencing (Eurofins and Genewiz) and purified with HiPure Plasmid Maxiprep kits (Invitrogen).

### Lentivirus production

Lentiviral particles were produced using the second-generation packaging system. Lentivirus encoding both LifeAct-RFP and EB3-YFP (cloned in a PLV-mCherry derived vector) were produced by co-transfection with the psPAX2 and pCMV-VSV-G helper plasmids (Addgene plasmids # 12260 and # 8454, gifts from Didier Trono and Bob Weinberg, respectively), into HEK293T cells (ATCC-CRL-3216) using the calcium phosphate transfection method. Viral particles were collected 48 h after transfection by ultra-speed centrifugation, prior to aliquoting and storage at −80°C.

### Primary hippocampal and cortical neuronal cultures

Mouse hippocampus and cortices were dissected from E18.5 embryos and digested in 0.25% trypsin in Hanks’ balanced salt solution (HBSS, Invitrogen, France) at 37°C for 15. min. After manual dissociation, cells were plated at a concentration of 5,000-15,000 cells/cm2 on 1 mg/ml poly-L-lysine-coated coverslips for fixed samples, or on ibidi glass bottom 60 µDishes for live imaging or plastic petri dishes for immunoblot analysis. Neurons were incubated 2 h in DMEM-10% horse serum and then changed to MACS neuro medium (Miltenyl Biotec) with B27 supplement (Invitrogen, France). One third of the medium was changed every 3–4 d up to 3 weeks in culture.

### Lentivirus infection

To perform dendritic spine quantification, a lentivirus containing eGFP at a multiplicity of infection of 5 was added to the cells at 7 DIV. Hippocampal neurons were incubated until 18 DIV at 37°C, 5% CO2 in a humidified incubator and then fixed with 4% paraformaldehyde in 4% sucrose-containing PBS for 20 min. To analyze microtubule invasion into the spines, mouse hippocampal neurons from wild type embryos were grown on 35 mm glass bottom live imaging dishes (ibidi) and infected at 7 DIV with a lentivirus containing LifeAct-RFP and EB3-YFP cDNA at a multiplicity of infection of 5.

### L-Dopa treatment

L-Dopa was incorporated into cultured hippocampal neurons as previously described (Dentesano et al., 2018; Zorgniotti et al., 2021). Briefly, the culture medium was substituted with Hank’s Balanced Salt Solution (HBSS; composition: 137 mM NaCl, 5 mM KCl, 0.8 mM MgSO4, 0.33 mM Na2HPO4, 0.44 mM KH2PO4, 0.25 mM CaCl2, 1 mM MgCl2, 0.15 mM Tris-HCl, pH 7.4), supplemented with freshly prepared 1 mM sodium butyrate. Additionally, 1 μM tolcapone (Sigma-Aldrich) and 50 μM carbidopa (Sigma-Aldrich) were added to the solution. The solution was then either supplemented with or without 0.4 mM L-Dopa (Sigma-Aldrich). The cells were then incubated for 1 hour at 37°C in a humidified atmosphere with 5% CO2.

### Immunofluorescence and antibodies

For immunocytochemistry, cells were fixed with 4% paraformaldehyde in 4% sucrose-containing PBS for 20 min and permeabilized with 0.2% Triton X-100/ PBS for 5 min. Fixed cells were then incubated with primary antibodies for 3 h in 0.1% PBS/Tween and then with Fluorophore-conjugated secondary antibodies for 1 h at room temperature. Primary antibodies were: mouse monoclonal anti PSD-95 (clone K28/43, NeuroMab UC Davis/NIH NeuroMab Facility. Item # 75-028) diluted 1:500, rabbit polyclonal anti Synaptophysin (SYP (H-93), Santa Cruz Biotechnology, Inc. Cat# sc-9116) diluted 1:500, rat monoclonal anti Tyr-tubulin (YL1/2) diluted 1:1000, rabbit polyclonal anti-deTyr tubulin diluted 1:1000, rabbit polyclonal anti-Δ2 tubulin diluted 1:1000 and mouse monoclonal antibody against α-tubulin (clone α3A1) diluted 1:1000 were previously described (Peris, Parato, Qu, Soleilhac, Lante, et al., 2022). Secondary antibodies were coupled to Alexa-488, to Alexa-565 or to Alexa-647 (Jackson Immuno-Research Laboratories).

### Imaging of dendritic spines

For cultured samples, hippocampal neurons from wild type, SVBP KO and TTL KO embryos were infected with eGFP containing lentivirus and fixed at DIV18 prior to mounting with DAKO antifade mounting reagent (Agilent S3023). Dendritic segments visualized by eGFP were obtained using an inverted microscope (Axio Observer, Zeiss) coupled to a spinning-disk confocal system (CSU-W1-T3, Yokogawa) and a LiveSR (super resolution) module connected to a wide-field electron-multiplying charge-coupled device (CCD) camera (ProEM+1024, Princeton Instrument). Serial optical sections with pixel dimensions of 0.104 × 0.104 μm were collected at 200 nm intervals for 10-15 stacks, using a × 63 oil-immersion objective (NA 1.46). Z stack images were shown as maximum projections. Dendritic spine analysis (spine counting and shape classification) was performed on the stacks using Neuronstudio as in (Peris, Parato, Qu, Soleilhac, Lante, et al., 2022). All spine measurements were performed in 3D from the z-stacks. The linear density was calculated by dividing the total number of spines present on assayed dendritic segments by the total length of the segments. At least two dendritic regions of interest were analyzed per cell from at least three independent embryos per culture in each experimental condition.

### Imaging of excitatory synapses

For immunodetection of excitatory synapses in eGFP expressing neurons, cells were fixed at 18 DIV and immunolabeled with PSD-95 and Synaptophysin antibodies. Fluorescent images were acquired with an inverted microscope (Axio Observer, Zeiss) coupled to a spinning-disk confocal system (CSU-W1-T3, Yokogawa) and a LiveSR (super resolution) module connected to a wide-field electron-multiplying charge-coupled device (CCD) camera (ProEM+1024, Princeton Instrument). Serial optical sections were acquired with 4 z-stack planes at 0.7 µm step size using Metamorph software. Images were enhanced for small structures with the LoG3D ImageJ plugin68 using a 2-pixel radius, and thresholded to create a mask for: eGFP (cell filler and contour marker), Synaptophysin (pre-synpatic compartment) and PSD-95 (post-synpatic compartment). The mask corresponding to the overlap of the three markers was superposed to the eGFP image. The synaptic puncta present in dendritic spines were manually counted. Mask creation and counting were done blind to the genotype and treatment.

### Live imaging of microtubule dynamics and dendritic spine invasion

LifeAct-RFP and EB3-YFP expressing hippocampal neurons (DIV 18) were incubated 1 h in HBSS with or without L-Dopa, as previously described. Initial conditioned MACs complete medium was added before transferring the cells to an inverted microscope (Axio Observer, Zeiss) coupled to a spinning-disk confocal system (CSU-W1-T3, Yokogawa) connected to a wide-field electron-multiplying charge-coupled device (CCD) camera (ProEM+1024, Princeton Instrument) with a 63×/1.46 oil objective and maintained at 37°C and 5% CO2. Movies of microtubule dynamics were acquired at 5 second/frame for 5 minutes with 4 z-stack planes at 0.7 µm step size by Metamorph software.

Maximum projections of movies were performed by ImageJ software, exported as Tiff files and analyzed in ImageJ. The percentage of the spines invaded was calculated as the number of spines invaded by microtubules during 5 min movie/total number of spines in the imaging field (Peris, Parato, Qu, Soleilhac, Lante, et al., 2022). To analyze microtubule dynamics, Kymographs were generated by drawing a region of the middle part of the dendrite. Parameters describing microtubule dynamics were defined as follows: catastrophe frequency: number of full tracks/total duration of growth; growth length: comet movement length in μm; comet lifetime: duration of growth; growth rate: growth length/comet lifetime (Peris, Parato, Qu, Soleilhac, Lante, et al., 2022).

### Analysis of spine structural plasticity

Dendritic spine morphology (stubby, mushroom, thin) of all protrusions invaded or not invaded by EB3 in the same imaging field before (0 h) and after L-Dopa treatment (1 h) were manually counted using ImageJ Software. Percentages of the same protrusions changing to pruned, thin, mushroom, or stubby spines were then calculated based on the total number of spines invaded or not invaded by EB3 in the same field. χ2 tests were performed on spine persistence or pruning in vehicle and L-Dopa treated neurons at 0 and 1 hours (Peris, Parato, Qu, Soleilhac, Lante, et al., 2022).

### Biochemical analysis of cultured mouse neurons

Cortical neurons (18 DIV) treated with HBSS (control) or with L-Dopa (1h) were collected, washed with phosphate-buffered saline medium at 37◦C and directly lysed in Laemmli buffer. L-Dopa incorporation into microtubules was calculated by semi-quantitative immunoblot analysis of tyrosinated, detyrosinated and Δ2 tubulin levels relative to total α-tubulin levels normalized to the corresponding modified mCherry α-tubulin standard, as in (Aillaud et al., 2016). We postulated that the sum of the analyzed modifications (Tyr, deTyr and Δ2 α-tubulin) represents 100% of α-tubulin in control neurons, and after L-Dopa treatment, the percentage of α-tubulin that is not Tyr, deTyr or Δ2 should be L-Dopa α-tubulin. Several neuronal cultures were used as indicated in the figure legends and for each sample, 2 independent blots were performed.

### Constructs and enzymes purification for TIRF and immunofluorescence studies

We used the human VASH1 (NP_055724)/human SVBP (NP_955374) complex fused to sfGFP tag, as described in (Ramirez-Rios et al., 2023). The VASH1 constructs used in this study were: VASH1 full-length (VASH1, residues 1–365) and the catalytically inactive VASH1 full length, (deadVASH1, residues 1–365, C169A)(Ramirez-Rios et al., 2023). Protein expression and purification of the various His-tagged VASH1–SVBP complexes were performed as previously described (Ramirez-Rios et al., 2023).

### Preparation of Tyr- and L-Dopa-tubulin and Taxol-stabilized microtubules

Preparation of brain tubulin and its labeling with either biotin or ATTO-565 fluorophore (ATTO-TEC Gmbh) were performed according to (Ramirez-Rios et al., 2023).

Aliquots of 2.5 mg of purified brain tubulin were incubated for 15 min at 30 °C in the presence of 2 U⋅mL-1 carboxypeptidase A (CPA) to remove the endogenous C-terminal tyrosine and increase the levels of detyrosinated tubulin that will act as substrate for tyrosine or L-Dopa incorporation (Lafanechère & Job, 2011). The CPA enzyme was subsequently inhibited by the addition of 20 mM dithiothreitol (DTT) and the tubulin samples were passed through a PD-10 desalting column filled with Sephadex G-25 resin. The eluted tubulin was centrifuged at 4 °C using a Vivaspin 500 concentrator (Sartorius) to achieve a final concentration of 10 mg⋅mL-1.

Parallel incorporation of L-Dopa or tyrosine into the purified detyrosinated tubulin was carried out using an incubation system composed of: 10 mg⋅mL-1 tubulin, 1 mM DTT, 2.5 mM ATP, 12.5 mM MgCl2, 100 mM KCl, 12 mM ascorbic acid, and 0.5 mM L-Dopa (Sigma-Aldrich) or 1 mM L-Tyrosine (Sigma-Aldrich), respectively (Rodriguez et al., 1975), in BRB80 buffer (80 mM Pipes-KOH at pH 6.8, 1 mM MgCl2, 1 mM EGTA). 0.25 mg⋅mL-1 of the recombinant TTL enzyme was added to the system and incubated for 30 min at 35 °C. After the incorporation of L-Dopa or tyrosine, the samples were cooled to 4 °C for 10 min and centrifuged at 200,000 x g for 10 min. The supernatants were collected and incubated for 1 h at 35 °C in presence of 30% glycerol, 1 mM GTP, and 5 mM MgCl2 to induce microtubule polymerization.

Microtubules enriched in Tyr- or L-Dopa-tubulin were centrifuged through a 60% glycerol cushion (v/v in BRB80) at 200,000 x g for 30 min at 35 °C. The pellet was resuspended in 15 μL of BRB80 buffer, and all tubulin proteins were stored in liquid nitrogen. Taxol-stabilized microtubules were prepared, as described in (Ramirez-Rios et al., 2023), by polymerizing 45 µM tubulin (composed of 65% of Tyr- or L-Dopa-tubulin, 30% biotinylated brain tubulin, and 5% ATTO-565–labeled brain tubulin) in BRB80 supplemented with 1 mM GTP. To allow the formation of microtubule, the mixture was incubated for 1 h at 35 °C. Subsequently, 100 μM Taxol was added and incubation was continued for 30 min. Tyr- or L-Dopa-microtubules were then centrifuged 10 min at 200,000 xg and resuspended in BRB80 supplemented with 10 μM Taxol.

### TIRF in vitro assay

Flow chambers for TIRF imaging assays were prepared as previously described (Ramirez-Rios et al., 2023). Subsequently, solution of microtubules enriched in either Tyr- or L-Dopa-tubulin was perfused into and incubated at room temperature for 5 – 7 min. Unbounded microtubules were removed by three washes of 100 µl of 1% BSA/10 µM Taxol in BRB80, supplemented with 10 μM Paclitaxel. Finally, 30 µl of a solution containing 25 pM of the catalytically inactive sfGFP–VASH1–SVBP complex (Ramirez-Rios et al., 2023) in BRB80 (supplemented with 82 µg/ml catalase, 580 µg/ml glucose oxidase, 1 mg/ml glucose, 4 mM DTT, 0.5 mM Taxol, 0.01% methylcellulose 1,500 cp) were perfused. The chamber was sealed, and images were recorded within the first 30 min following addition of the assay-mix solution on an inverted microscope (Eclipse Ti, Nikon). The microscope was equipped with a Perfect Focused System, a CFI Apochromat TIRF 100×/1.49 N.A. oil immersion objective (Nikon), a warm stage controller (Linkam Scientific) and a Technicoplast chamber to maintain the temperature, an objective heater (OkoLab), an iLas2 TIRF system (Roper Scientific), and a sCMOS camera (Prime95B, Photometrics) controlled by MetaMorph software (version 7.10.3, Molecular Devices). For dual view imaging, an OptoSplit II bypass system (Cairn Research) was used as image splitter and illumination was provided by 488- and 561-nm lasers (150 and 50 mW, respectively). Temperature was maintained at 35°C for all imaging purposes. Acquisition rate was one frame each 50 ms exposure (in streaming acquisition) during 45 s.

### In vitro assay of detyrosination activity using immunofluorescence

VASH1-SVBP activity was measured as in (Ramirez-Rios et al., 2023). Immunofluorescence in vitro assays were made using exactly the same perfusion chambers and TIRF experiments conditions as above. After addition of the assay-mix solution, incubation was done at 37°C for 30 min followed by three washes with 10 µM Taxol and 1% BSA in BRB80 (wash buffer). The incubation with primary antibodies (rat anti-tyrosinated tubulin (YL1/2 = anti-Tyr, 1:6,000) and rabbit anti-detyrosinated tubulin (anti-deTyr, 1:1,000)) was done for 15 min, followed by three washes with 100 µl of wash buffer. Subsequently, secondary antibodies (anti-rat coupled to Alexa Fluor 488 (Jackson ImmunoResearch, 712-545-153) and anti-rabbit coupled to Cyanine 3 (711-165-152; Jackson ImmunoResearch), both diluted to 1:500) were added and incubated for 15 min, followed by three washes with 100 µl of wash buffer. Images were obtained using a LEICA DMI600/ROPER microscope controlled by Metamorph Video software using the same illumination conditions. For each condition at least three independent experiments with different protein preparations were done.

### Data analysis

Immunofluorescence and binding tracking of single VASH1–SVBP molecules for estimation of activity and binding parameters were measured using FIJI software and a homemade plugin KymoTool (Ramirez-Rios et al., 2016), as previously described in (Ramirez-Rios et al., 2023).

### Statistical analysis

All data are presented as mean ± SEM. Statistical significance of differences between conditions was calculated with Prism 7.0 (GraphPad Software), using tests as indicated in each figure. Mean differences were considered significant at * p < 0.05; ** p < 0.01; *** p < 0.001 and **** p < 0.0001.

## RESULTS

### L-Dopa incorporation into microtubules reduces dendritic spine density in wild type hippocampal neurons

Previous studies have demonstrated that L-Dopa, a tyrosine analog commonly used as standard treatment in Parkinson’s disease, can be incorporated at the C-terminal tail of α-tubulin by the enzyme TTL and that L-Dopa tubulin can polymerize into microtubules (Dentesano et al., 2018; Rodriguez et al., 1975; Zorgniotti et al., 2021)(Fig 1 A).

**Figure 1.**
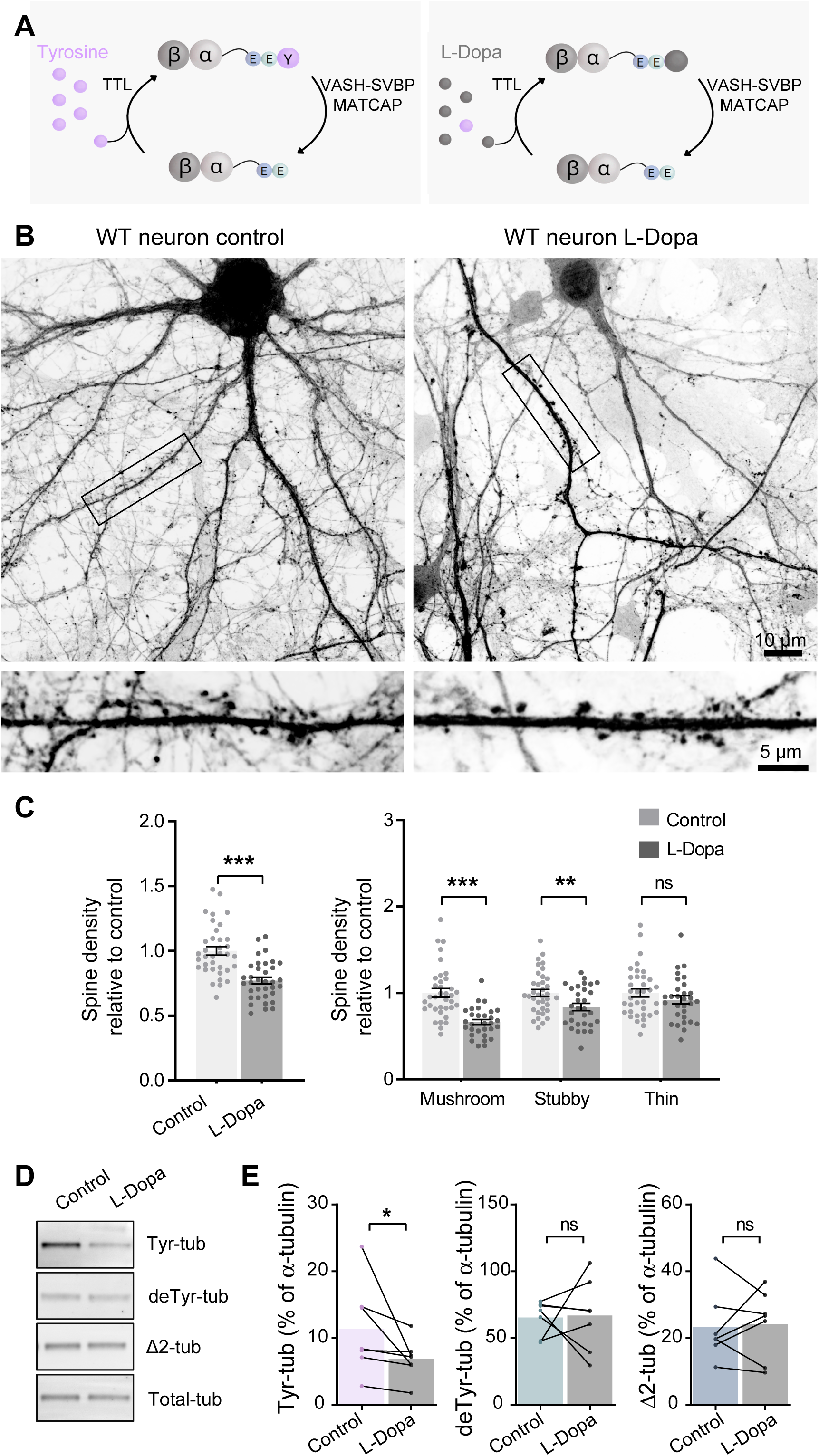
L-Dopa incorporation into microtubules reduces dendritic spine density in wild type hippocampal neurons. **(A)** Schematic representation of αβ tubulin dimers with their C-terminal tails. Under physiological condition (left) tubulin carboxypeptidases (VASH1/2-SVBP, and MATCAP) remove the C-terminal tyrosine residue (Y, in violet) of α-tubulin to generate detyrosinated tubulin. Detyrosinated tubulin can be re-tyrosinated by the tubulin tyrosine ligase (TTL) that adds a tyrosine residue at the glutamate C-terminal (E, in green). In the presence of L-Dopa (right, in gray), this amino acid is incorporated into the C-terminal of detyrosinated tubulin by the TTL enzyme, replacing the tyrosine residue in the detyrosination/tyrosination cycle. **(B)** Representative confocal images of eGFP expressing wild-type (WT) hippocampal neurons (18 DIV) treated with 0.4 mM L-Dopa or the vehicle (Control) for 1 h. Scale bar: 10 µm **(C)** Total dendritic spine density, or that of each different morphological type of spines, is represented for wild type neurons incubated with L-Dopa over 1 h. Spine density values were normalized to the mean of the control cells. Graphs represent mean ± SEM; n = 36 control and n = 35 L-Dopa-treated neurons from three independent cultures. Mann-Whitney test, **P < 0.01; ***P < 0.001 and ns = not significant. **(D)** Representatives immunoblot of protein extracts from cortical neurons (18 DIV) treated with 0.4 mM L-Dopa or the vehicle (Control) for 1h showing tyrosinated (Tyr-tub), detyrosinated (deTyr-tub), Δ2 (Δ2-tub) and total α tubulin (Total-tub) levels. **(E)** The mean content of the different forms of α-tubulin (tyrosinated, detyrosinated and Δ2 tubulin) present in the samples was estimated after normalization to total α-tubulin levels and antibody sensitivity (determined by the co-analysis of extracts from HEK293T cells transfected with various m-Cherry α-tubulin variants (not showed), described in methods section. Wilcoxon test *P < 0.05; ns = not significant.

Mature wild type hippocampal neurons treated with L-Dopa (0.4 mM, 1h) showed reduced dendritic spine density (Fig 1 B, C, 1.00 ± 0.03 spines/µm and 0.77 ± 0.02 spines/µm for control and L-Dopa treated, respectively), affecting mainly the mature forms of dendritic spines (34 ± 10 % and 16 ± 9 % of reduction in mushrooms and stubby spines, respectively) indicating a deleterious consequence of L-Dopa incorporation into tubulin and the formation of L-Dopa-microtubules on the maintenance of the dendritic spines. The global morphology of the spines was unaffected after L-Dopa treatment (Fig S1) suggesting that L-Dopa treatment induced spine pruning and/or changes from one spine subpopulation to others.

The analysis of the tyrosination state the microtubular network on these neurons demonstrated that L-Dopa treatment induced almost 40% of reduction in the levels of Tyr-tubulin (Fig 1 D-E, 11 ± 3 % vs 7 ± 1 % of total α-tubulin for control and L-Dopa treated neurons, respectively) with no significant change in the levels of detyrosinated and Δ2-tubulin (65 ± 5 vs 67 ± 10 % and 23 ± 4 vs 24 ± 4 % of total α-tubulin for control and L-Dopa treated neurons, respectively), indicating the presence of a new tubulin pool that is recognized by total α-tubulin antibody, but is neither tyrosinated, detyrosinated, nor Δ2-tubulin. As there is not an available L-Dopa-tubulin antibody, we can speculate that this new tubulin pool that appears after L-Dopa treatment is L-Dopa-α-tubulin, and correspond to approximately 40% of the Tyr-tubulin present in not treated wild type neurons (Fig 1 E), coinciding with what was previously reported by (Zorgniotti et al., 2021).

### L-Dopa incorporation into microtubules perturbs VASH1-SVBP carboxypeptidase activity by altering microtubule interaction

We analyzed the release of L-Dopa from microtubules by evaluating the activity of the VASH1-SVBP complex on L-Dopa-tubulin enriched microtubules by in vitro assays. To generate Tyr- or L-Dopa-modified microtubules, we first induced α-tubulin detyrosination by incubating purified bovine brain tubulin with carboxypeptidase A (CPA). After CPA inactivation, soluble deTyr-tubulin was incubated with the recombinant TTL enzyme and either L-Tyrosine or L-Dopa (Fig 2 A). As expected, we observed a significant increase in Tyr-tubulin levels when deTyr-tubulin was incubated with TTL and L-Tyrosine, and no change when incubated with TTL and L-Dopa (Fig 2 B, from 0.11 ± 0.07 a.u. to 1.3 ± 0.3 a.u. for TTL + Tyr and from 0.11 ± 0.07 a.u. to 0.04 ± 0.04 a.u. for TTL + L-Dopa). We observed an important reduction of deTyr-tubulin levels in both conditions (from 4.0 ± 0.4 a.u. to 0.4 ± 0.2 a.u. for TTL + Tyr and from 4.0 ± 0.4 a.u. to 0.4 ± 0.1 a.u. for TTL + L-Dopa) with no major changes in the levels of Δ2-tubulin (from 3 ± 1 a.u. to 3 ± 1 a.u. for TTL + Tyr and from 3 ± 1 to 2 ± 1 a.u. for TTL + L-Dopa, Fig 2 B). The reduction of deTyr-tubulin levels with no change in Tyr- and Δ2-tubulin provided first-time evidence of L-Dopa incorporation into tubulin by recombinant TTL in purified systems.

**Figure 2.**
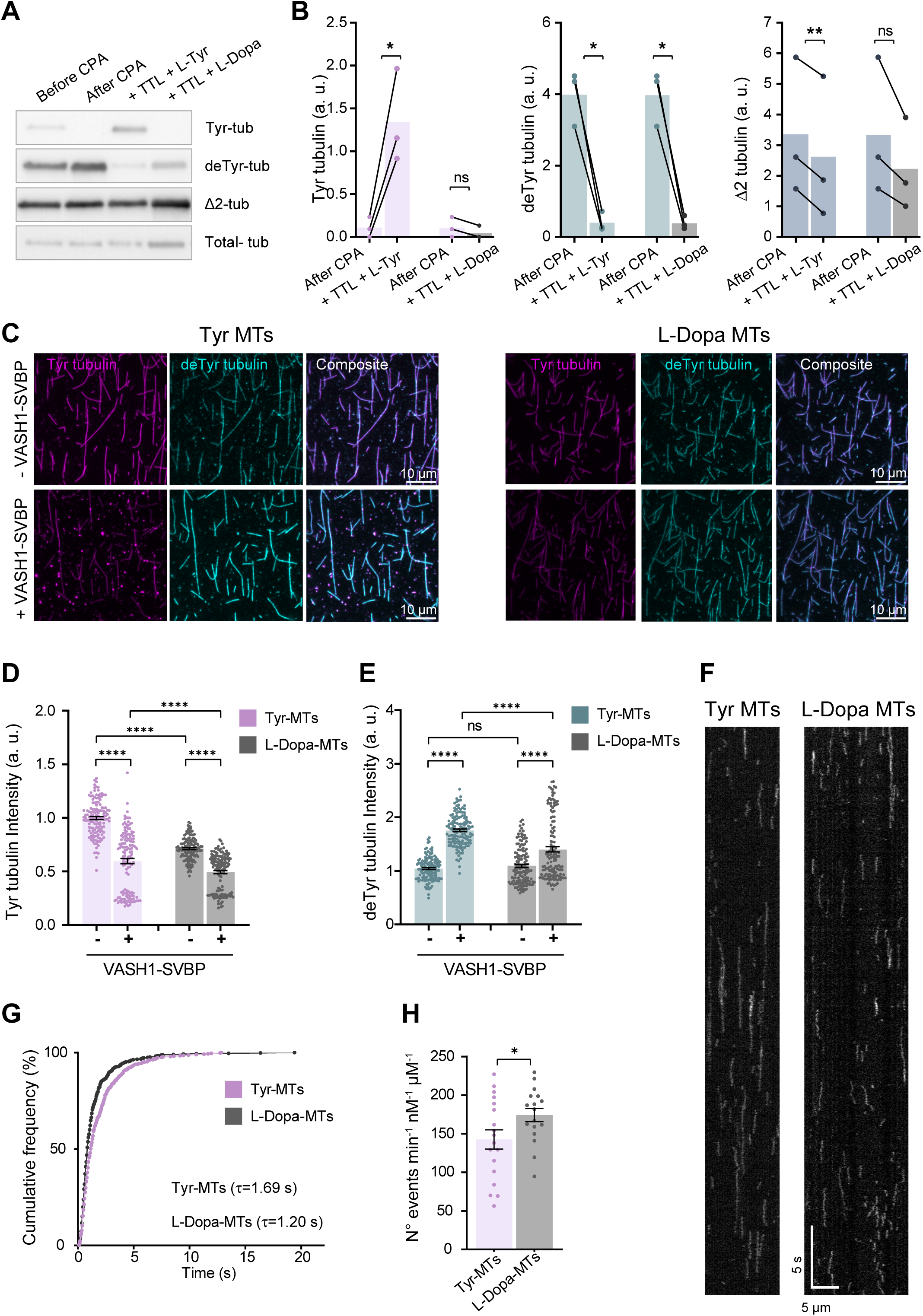
L-Dopa incorporation into microtubules perturbs VASH1-SVBP carboxypeptidase activity by altering microtubule interaction. **(A)** Representatives immunoblot of purified bovine brain tubulin fractions showing tyrosinated (Tyr-tub), detyrosinated (deTyr-tub), Δ2 (Δ2-tub) and total α tubulin (Total-tub) levels. The levels of the modified tubulins were analyzed in the initial tubulin preparation (Before CPA), after the incubation with carboxypeptidase A (After CPA) and following the incubation with the TTL enzyme in the presence of L-Tyrosine (+TTL +Tyr) or L-Dopa (+TTL +Dopa). **(B)** Immunoblot quantification of modified tubulins (tyrosinated, detyrosinated and Δ2 tubulin) present in the different samples. The levels of the modified tubulins were estimated after normalization to the corresponding total α-tubulin levels. The graphs show the variation between the initial tubulin levels (After CPA) and after the incorporation of tyrosine (+TTL +Tyr) or L-Dopa (+TTL +Dopa). Paired t-test **P < 0.01; *P < 0.05; ns = not significant. **(C)** Representative images of microtubules enriched in tyrosinated (Tyr MTs, left panel) or L-Dopa (L-Dopa MTs, right panel) tubulin. For each type of microtubules, images of tyrosinated (Tyr, magenta) and detyrosinated (deTyr, cyan) α-tubulin pools after 30 min of incubation in the absence (-VASH1-SVBP) or presence (+VASH1-SVBP) of the VASH1-SVBP complex are shown. Scale bar 10 µm. **(D - E)** Analysis of tyrosinated tubulin **(D)** and detyrosinated tubulin **(E)** signal intensities. Each point represents an individual microtubule. The fluorescence intensity values were normalized to the mean of the tyrosinated microtubules in the absence of enzyme. Data represent mean ± SEM; n = 138 microtubules per condition, 3 independent experiments. Kruskal-Wallis test; **** p < 0.0001. **(F)** Representative kymographs of single molecules of catalytically inactive sfGFP-tagged VASH1–SVBP at a concentration of 25 pM bound to Taxol-stabilized microtubules enriched in tyrosinated (Tyr MTs) or L-Dopa (L-Dopa MTs) tubulin. Scale bars: horizontal, 5 µm; vertical, 5 s. **(G)** Cumulative frequency of the residence times measured in TIRF movies taken during the 30 min following addition of enzyme complexes to tyrosinated (Tyr MTs) or L-Dopa (L-Dopa MTs) microtubules. The mean residence time (τ) is obtained by fitting the curve with a mono-exponential function, described in methods section(Ramirez-Rios et al., 2023)).**(H)**. Analysis of binding frequency. Number of binding events of the deadVASH1-SVBP enzyme complex to tyrosinated (Tyr MTs) or L-Dopa (L-Dopa MTs) microtubules. Each point represents an individual microtubule. Data represent mean ± SEM; n = 18 Tyr MTs and n = 17 L-DopaMTs. Unpaired t-test; *P < 0.05.

We next analyzed both the detyrosinating activity as well as the interaction with microtubules of VASH1–SVBP complex (the major detyrosinating enzyme in brain) using immunofluorescence and TIRF microscopy, respectively. We immobilized Taxol-stabilized microtubules enriched in tyrosinated or L-Dopa tubulin on the surface of TIRF chambers as previously described (Ramirez-Rios et al., 2023) and we tested the activity of VASH1–SVBP in vitro using immunofluorescence (Fig 2 C).

In control conditions, without purified VASH1-SVBP, the Tyr-tubulin content of Tyr-microtubules was significantly higher than the Tyr-tubulin content of L-Dopa-microtubules (1.00 ± 0.01 a.u. vs 0.71 ± 0.01 a.u. for Tyr- and L-Dopa-microtubules, respectively, Fig 2 D) while the deTyr-tubulin content of both was similar (1.04 ± 0.02 a.u. and 1.40 ± 0.05 a.u. for Tyr- and L-Dopa-microtubules, respectively, Fig 2 E). After incubation with VASH1-SVBP, Tyr-tubulin content was significantly reduced (40 ± 5 %) in Tyr-microtubules whereas the reduction was only of 31 ± 4 % in L-Dopa-microtubules (Fig 2 D). As expected, VASH1-SVBP significantly increased deTyr-tubulin content in Tyr-microtubules (69 ± 3 %) whereas the increase was only of 28 ± 6 % on L-Dopa-microtubules (Fig 2 E). The significantly larger detyrosination of Tyr-microtubules compared to L-Dopa-microtubules indicated a reduced carboxypeptidase activity of VASH1-SVBP on L-Dopa-microtubules.

In order to understand the molecular mechanisms involved in the distinct detyrosination activity on L-Dopa-microtubules, we examined the microtubule-interacting behavior of the catalytically inactive enzyme complexes containing a Cys169Ala mutation in VASH1 (deadVASH1-SVBP,(Ramirez-Rios et al., 2023)). Interestingly, as shown in representative kymographs, the binding behavior of the enzyme complex on the two types of microtubules was very different (Fig. 2 F). We observed that deadVASH1-SVBP bound more frequently (143 ± 12 events/min.µm.nM vs 174 ± 9 events/min.µm.nM for Tyr- and Dopa-microtubules, respectively; Fig 2 G) and for much shorter times on L-Dopa-microtubules (τ = 1.7 s vs τ = 1.2 s for Tyr- and L-Dopa microtubules, respectively, Fig 2 H). This phenomenon was even more significant when the microtubules were exposed to a higher concentration of the deadV1-SVBP complex (Fig S2). We found a two-fold increase in binding frequency of the enzyme complex to L-Dopa-microtubules (72 ± 5 events/min.µm.nM vs 130 ±7 events/min.µm.nM for Tyr- and L-Dopa-microtubules, respectively) and a 30 % reduction in the residence time (τ = 2.2 s vs τ = 1.5 s, respectively).

Together, these results clearly highlight the different carboxypeptidase activity and binding behavior of the VASH1–SVBP enzyme on Tyr- and L-Dopa-microtubules. These anomalies could potentially contribute to a cumulative impact on L-Dopa-microtubules and exacerbate their atypical behavior.

### L-Dopa treatment does not reduce dendritic spine density in TTL KO or SVBP KO hippocampal neurons

To validate that dendritic spine defects observed in wild type neurons after L-Dopa treatment were due to the presence of L-Dopa-microtubules, we decided to use two cellular models in which, due to the absence of key enzymes of the detyrosination/tyrosination cycle, the post-translational incorporation of L-Dopa into tubulin does not occur: hippocampal cultured neurons from TTL and SVBP KO embryos (Fig 3 A, B).

**Figure 3.**
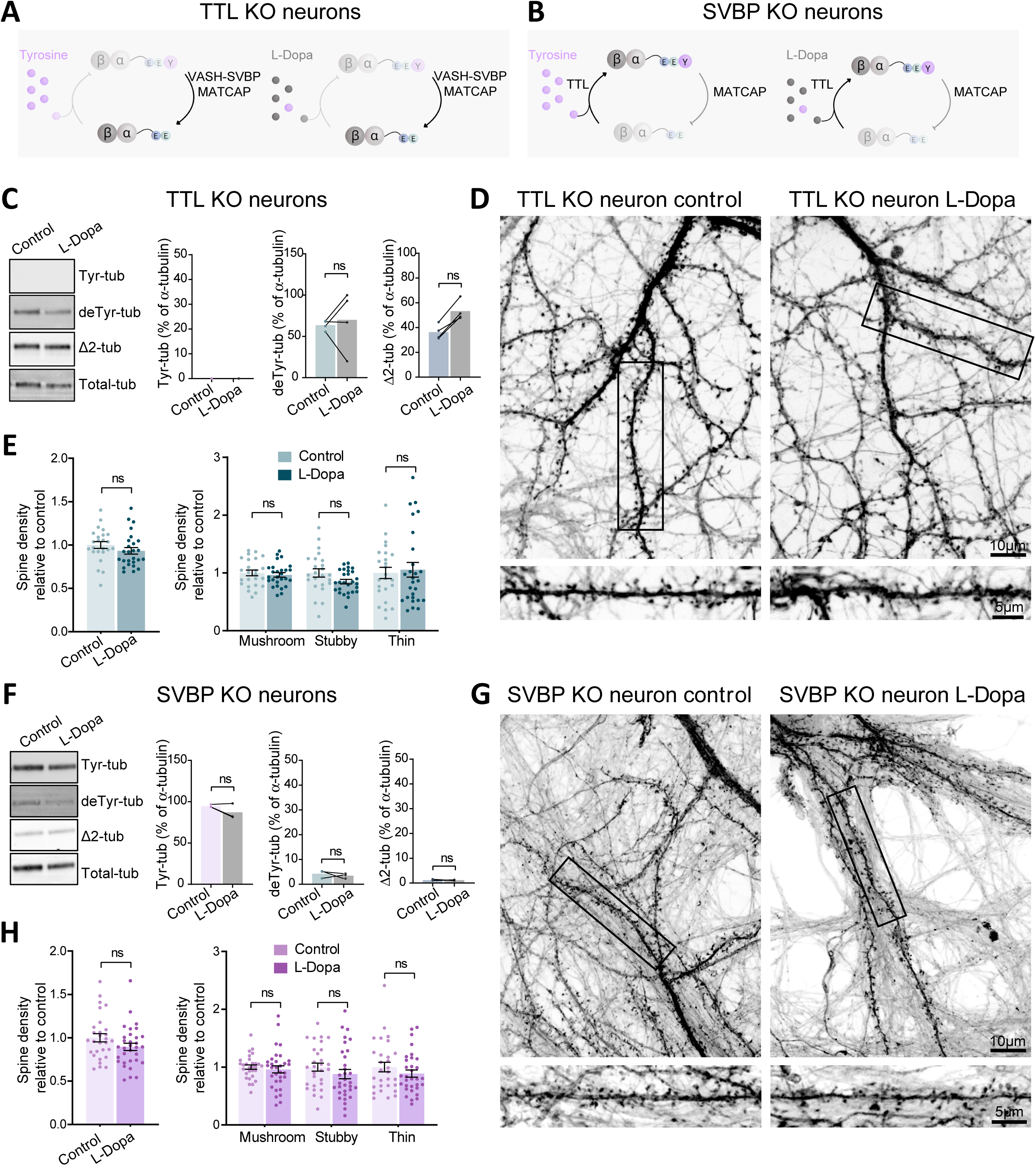
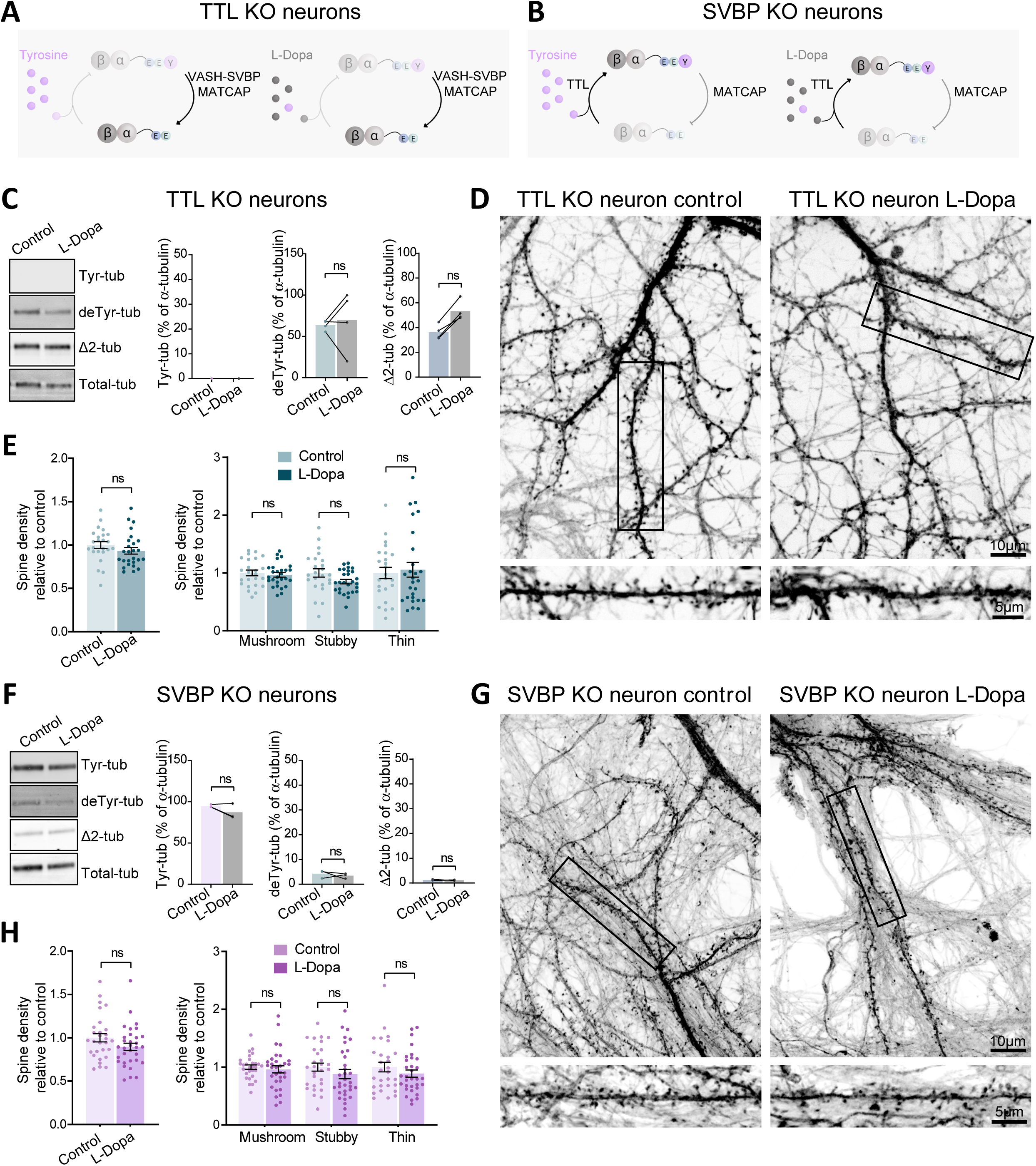
L-Dopa treatment does not reduce dendritic spine density in TTL KO or SVBP KO hippocampal neurons. **(A - B)** Schematic representation of the detyrosination/tyrosination cycle in the presence of tyrosine (violet) or L-Dopa (gray) in TTL knockout (KO) **(A)** or SVBP KO neurons **(B)**. **(C - F)** Representative immunoblot of protein extracts from TTL KO **(C)** and SVBP KO **(F)** cortical neurons (18 DIV) treated with 0.4 mM L-Dopa or the vehicle (Control) for 1h, showing tyrosinated (Tyr-tub), detyrosinated (deTyr-tub), Δ2 (Δ2-tub) and α tubulin (α-tub) levels. The content of the different forms of α-tubulin (tyrosinated, detyrosinated and Δ2) present in the samples was estimated after normalization to total α-tubulin levels and antibody sensitivity (determined by the co-analysis of extracts from HEK293T cells transfected with various mCherry α-tubulin variants, as described in methods section). Wilcoxon test; ns = not significant. **(D - H)** Representative confocal images of eGFP expressing TTL KO **(D)** and SVBP KO **(G)** hippocampal neurons (18 DIV) treated with 0.4 mM L-Dopa or the vehicle (Control). Scale bar: 10 µm. Total dendritic spine density, or that of each different morphological type of spines, is represented for TTL KO **(E)** and SVBP KO **(H)** neurons. Spine density values were normalized to the mean of the control cells. Graphs represent mean ± SEM; n = 24 control and n = 27 L-Dopa-treated TTL KO neurons and n = 29 control and n = 32 L-Dopa-treated SVBP KO neurons from three independent cultures. Mann-Whitney test, ns = not significant.

In TTL KO neurons, neither tyrosine nor L-Dopa can be incorporated into the C-terminus of α-tubulin due to the absence of the ligase enzyme (Fig 3 A). In these neurons, the significant increase in deTyr- and Δ2-tubulin levels (Erck et al., 2005) was unaffected by the treatment with L-Dopa (64 ± 3 % vs 70 ± 18 % for deTyr-tubulin; 36 ± 3 % vs 53 ± 4 % for Δ2-tubulin in control and L-Dopa treated neurons, respectively; Fig 3 C). L-Dopa treatment did not perturb dendritic spine density (1.00 ± 0.04 spines/µm vs 0.93 ± 0.04 spines/µm for control and L-Dopa treated neurons, respectively; Fig 3 D, E) or dendritic spine subpopulation (1.00 ± 0.05 spines/µm vs 0.97 ± 0.04 spines/µm, 1.00 ± 0.07 spines/µm vs 0.85 ± 0.04 spines/µm, 1.0 ± 0.1 spines/µm vs 1.0 ± 0.1 spines/µm for mushroom, stubby and thin spines in control and L-Dopa treated neurons; Fig 3 E) in TTL KO cultured hippocampal neurons, indicating that the post-translational incorporation of L-Dopa into tubulin was necessary for the observed effect on dendritic spines.

As L-Dopa needs deTyr-tubulin to be incorporated in, we also analyzed the microtubular network of SVBP KO cultured hippocampal neurons. It has been shown that the absence of the chaperone SVBP results in a complete absence of detyrosinase activity for VASH1/2 enzymes, and a drastic reduction of deTyr-tubulin levels in cells (Aillaud et al., 2017; Martínez-Hernández et al., 2022; Nieuwenhuis et al., 2017; Pagnamenta et al., 2019; Wang et al., 2019) (Fig 3 B, F). As expected, L-Dopa treatment did not significantly modify the levels of Tyr-, deTyr- or Δ2-tubulin in SVBP KO cultured neurons (95 ± 1 % vs 87 ± 5%; 4.2 ± 0.9 % vs 3.4 ± 0.6 % and 1.2 ± 0.2 % vs 1.15 ± 0.09 % for Tyr-deTyr and Δ2-tubulin in control and L-Dopa treated neurons, respectively; Fig 3 F), most likely due to a reduction in the substrate for the TTL enzyme, which prevents the incorporation of L-Dopa into tubulin. Consequently, L-Dopa treatment did not perturb dendritic spine density nor dendritic spine subpopulation in SVBP KO cultured hippocampal neurons (1.00 ± 0.05 spines/µm vs 0.89 ± 0.04 spines/µm for control and L-Dopa treated neurons, respectively; 1.00 ± 0.04 spines/µm vs 0.96 ± 0.06 spines/µm, 1.00 ± 0.07 spines/µm vs 0.88 ± 0.08 spines/µm, and 1.00 ± 0.08 spines/µm vs 0.89 ± 0.06 spines/µm for mushroom, stubby and thin spines in control and L-Dopa treated neurons, respectively; Fig 3 G, H).

These results demonstrated that the impaired L-Dopa incorporation into microtubules in TTL KO and SVBP KO neurons prevented the L-Dopa-dependent reduction of dendritic spines, suggesting that the effect on synaptotoxicity is driven by abnormal L-Dopa microtubules.

L-Dopa incorporation into microtubules reduced excitatory synapses To determine whether changes in morphologically defined dendritic spines corresponded to synaptic alterations, we immuno-labeled excitatory synapses in GFP expressing cultured hippocampal neurons (Peris et al., 2018). Neurons were transduced with lentiviral vectors to express soluble GFP, treated with L-Dopa and fixed at DIV18. Immunostaining with anti PSD-95 and anti synaptophysin antibodies (a post-synaptic and a pre-synaptic marker, respectively) allowed us to create a mask with the triple labeling (GFP, PSD-95 and synaptophysin). Fluorescent puncta containing both pre and post-synaptic markers were used to detect and count synapses formed on the spines of transduced cells, as shown in Fig. 4 A-C. Wild type GFP-expressing neurons displayed a reduced percentage of spines containing excitatory synapses (73 ± 2 % of spines vs 66 ± 2 % of spines in control and L-Dopa treated neurons, respectively; Fig. 4 A, D), whereas TTL KO and SVBP KO GFP-expressing neurons showed no difference in the percentage of spines containing excitatory synapses after L-Dopa treatment (71 ± 2 % of spines vs 69 ± 2 % of spines in TTL KO neurons and 66 ± 2 % of spines vs 65 ± 2 % of spines in SVBP KO neurons in control and L-Dopa treated cells, respectively; Fig. 4 D). Thus, in wild type cultured hippocampal neurons, L-Dopa treatment not only decreased dendritic spine density, but also the percentage of excitatory synapses in the remaining spines. These results suggest that L-Dopa treatment may induce a cumulative defect on the synaptic compartment that would amplify the gravity of the synaptic phenotype.

**Figure 4.**
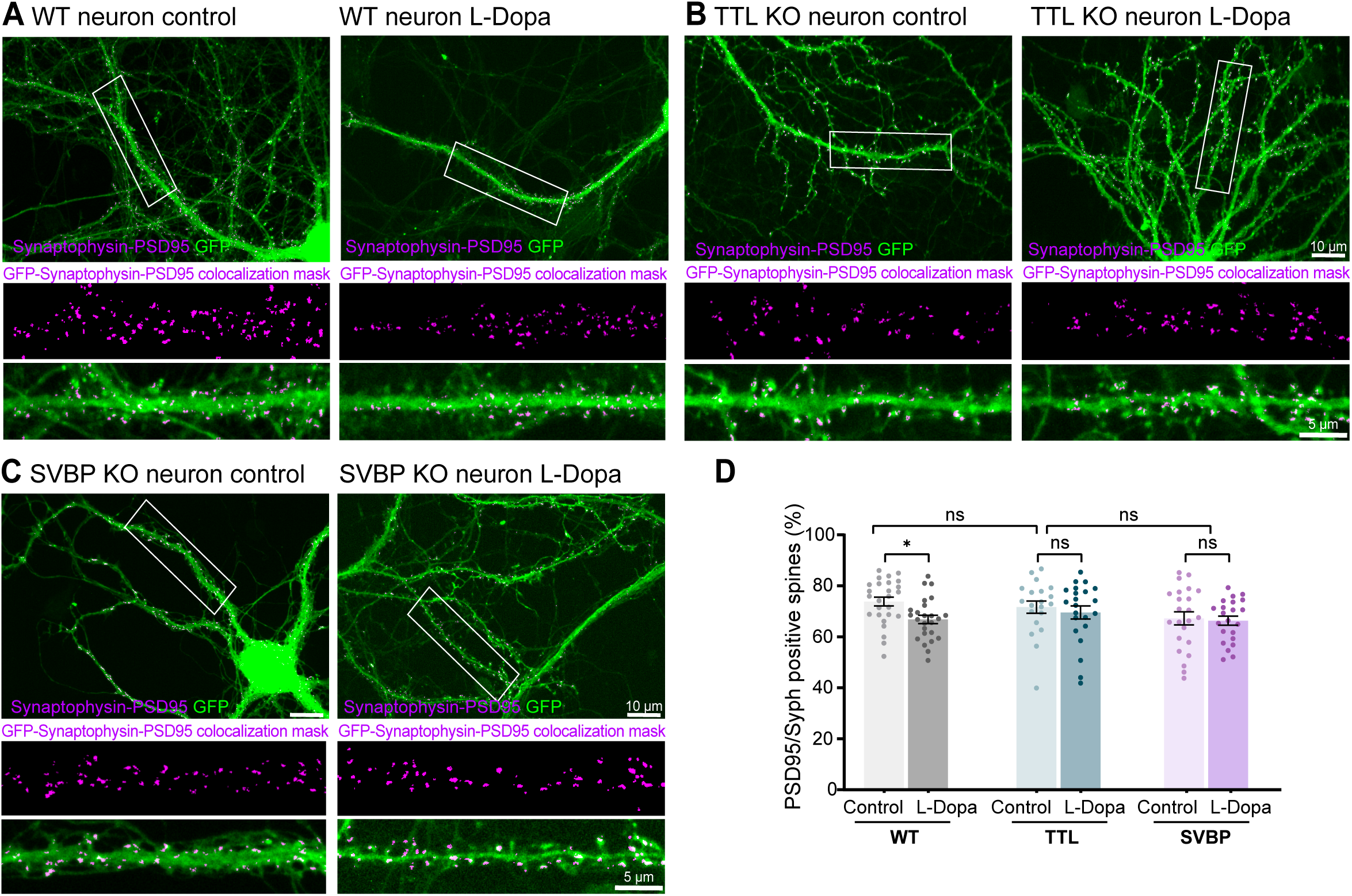
L-Dopa incorporation into microtubules reduces the number of excitatory synapses in wild type hippocampal neurons. **(A - C)** Representative confocal images of control and L-Dopa treated hippocampal wild-type (WT) **(A)**, TTL KO **(B)**, and SVBP KO **(C)** neurons (DIV18) expressing soluble GFP (green), including magnified views of the indicated dendritic segments. Images are superposed with a mask (magenta) corresponding to pixels that simultaneously represent PSD-95 (post-synaptic marker), Synaptophysin (pre-synaptic marker) and GFP label. Scale bar: 10 µm and 5 µm in the complete image and the magnified view, respectively. **(D)** Quantification of the percentage of dendritic spines containing fluorescent puncta of both pre- and post-synaptic markers in GFP expressing neurons. Data represent mean ± SEM; n = 20, n = 22 control and L-Dopa-treated WT neurons; n = 23, n = 22 control and L-Dopa-treated TTL KO neurons; and n = 26, n = 25 control and L-Dopa-treated SVBP KO neurons, respectively; from at least three different embryos and neuronal cultures. Kruskal-Wallis test; *P < 0.05; ns = not significant.

### L-Dopa incorporation modifies the microtubule dynamics and their ability to invade dendritic spines

Based on our findings in wild type neurons, where changes in dendritic spines and excitatory synapses relied on L-Dopa incorporation into tubulin, we subsequently investigated microtubule dynamics in the dendritic shaft. We transiently expressed the microtubule plus-end binding protein EB3-YFP to track the dynamic behavior of microtubule plus ends (Fig. 5 A). We found that in L-Dopa treated neurons, while comet growth length and speed were unchanged (4.4 ± 0.2 µm vs 4.0 ± 0.2 µm, and 9.2 ± 0.6 µm/min vs 9.1 ± 0.5 µm/min, for control and L-Dopa treated neurons, respectively; Fig. 5 B), catastrophe frequency was significantly increased with a corresponding decrease in comet lifetime compared to controls (1.83 ± 0.07 vs 2.05 ± 0.05, and 0.57 ± 0.02 min vs 0.50 ± 0.01 min for control and L-Dopa treated neurons, respectively; Fig. 5 B). These observations strongly suggest that the presence of L-Dopa on neuronal microtubules alters their dynamics.

**Figure 5.**
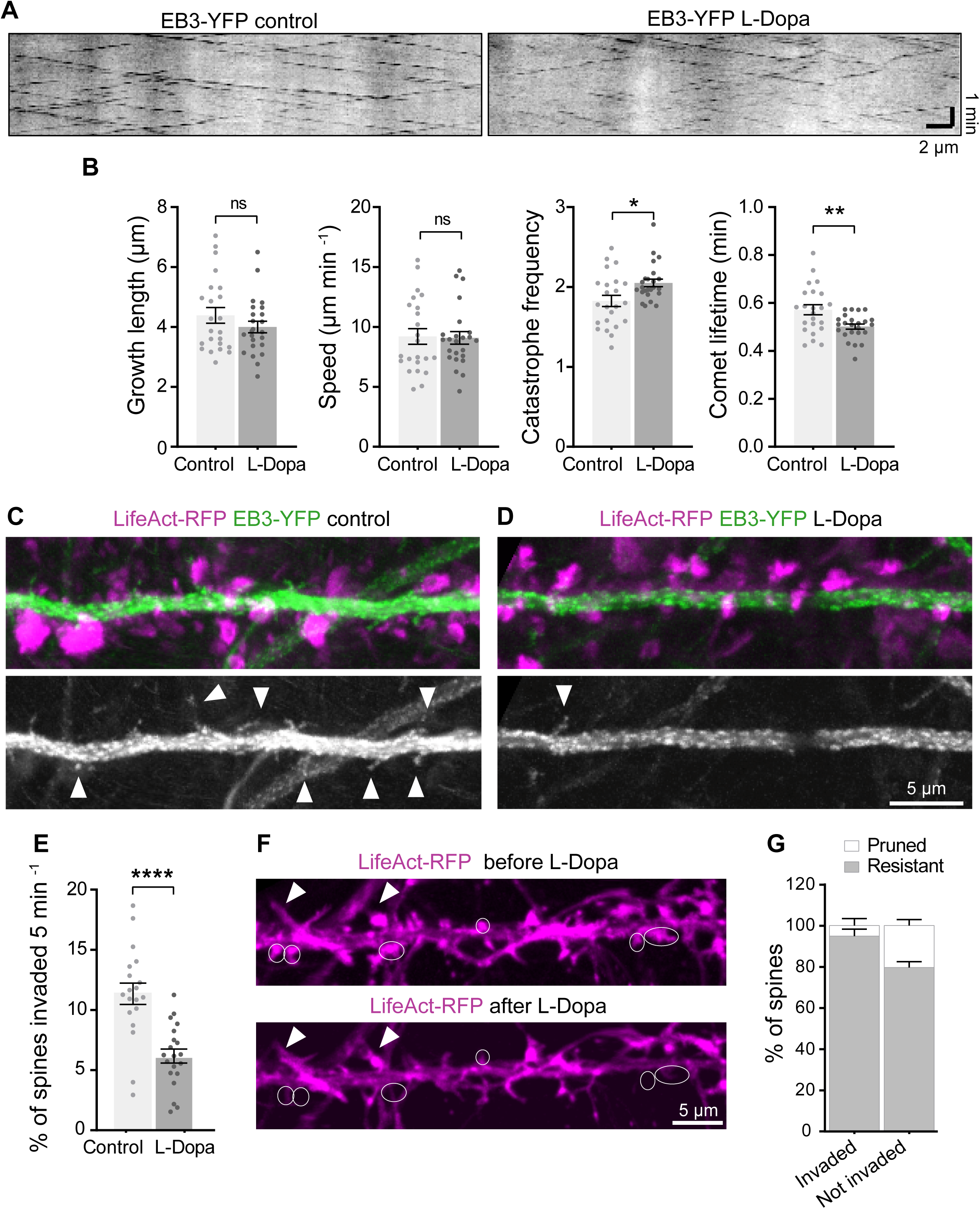
L-Dopa incorporation into microtubules modifies the MT dynamics and their ability to invade dendritic spines. **(A)** Representative kymographs of EB3-YFP comets corresponding to the plus ends of growing microtubules in a dendritic segment of 18 DIV wild type (wild type) hippocampal neurons either untreated (Control, left) or treated with 0.4 mM L-Dopa (L-Dopa, right). Scale bar: horizontal, 2 µm; vertical, 1 min. **(B)** Quantification of microtubule dynamics parameters including growth length, speed, catastrophe frequency and comet lifetime. Data represent mean ± SEM; n = 23, n = 24 control and L-Dopa-treated neurons, respectively from at least three different embryos and neuronal cultures. Mann-Whitney test, *P < 0.05; **P < 0.01 and ns = not significant. **(C - D)** Representative maximal intensity projection of time-lapse confocal images showing dendritic segments of WT hippocampal neurons (DIV 18) expressing both LifeAct-RFP (magenta) and EB3-YFP (green at top; gray at bottom) either untreated (Control) **(C)** or treated with 0.4 mM L-Dopa (L-Dopa) **(D)**. In the bottom panels, the white arrows indicate microtubules invading dendritic spines. Scale bar: 5 µm. **(E)** Percentage of spines invaded by microtubules in neurons treated either with the vehicle (Control) or 0.4 mM L-Dopa (+Dopa). Data represent mean ± SEM; n = 19, n = 20 control and L-Dopa-treated neurons respectively, from at least three different embryos and neuronal cultures. Unpaired t-test, ****P < 0.0001. **(F)** Representative images showing dendritic segments of a neuron (DIV18) expressing LifeAct-RFP (magenta) and EB3-YFP (not shown) before (top) and after (bottom) L-Dopa treatment. The white arrows point out the spines that were invaded by microtubules before L-Dopa treatment and the circular ROIs indicate the dendritic spines that will prune after L-Dopa treatment (bottom panel). **(G)** Total percentage of spines pruned (white) or resistant (gray) after L-Dopa treatment. Graph represents the mean percentage of microtubule-invaded (left) and non-invaded spines (right) for either fate; n = 6 cells from two independent experiments. Two-way ANOVA, ****P < 0.0001 (resistance vs pruning 92.82 < 0.0001 ** Yes).

We previously demonstrated that abnormal synaptic microtubule dynamics, caused by a disrupted tubulin detyrosination/tyrosination cycle, reduced the number of microtubules entering into dendritic spines, leading to a significant loss of synapses (Peris, Parato, Qu, Soleilhac, Lante, et al., 2022). We hypothesized that abnormal L-Dopa-microtubules dynamics might similarly affect spine invasion, potentially explaining the reduction in dendritic spines observed after L-Dopa treatment. We analyzed microtubule invasions into individual spines of wild type cultured hippocampal neurons treated or not with L-Dopa. In order to simultaneously label actin enriched in dendritic spines with LifeAct-RFP and microtubule plus-end with EB3-YFP (Fig 5 C, D) neurons were, infected with a lentivirus containing a home-made construction of LifeAct-RFP-IRES-EB3-YFP. L-Dopa treatment drastically reduced the percentage of spines invaded by microtubules (11 ± 4 % vs 6 ± 3 % of invaded spines in 5 min in control and L-Dopa treated neurons, respectively; Fig 4 C - E, white arrowheads showing invaded spines) indicating that abnormal dynamic behavior of L-Dopa-microtubules has a major impact on synaptic microtubules invasion. Next, we tracked and quantified the fate of the same spines, whether invaded or not by microtubules, before and after L-Dopa treatment (Fig. F, G). L-Dopa treatment had a minor effect on the pruning of microtubule-invaded spines (5 ± 3 % of spines), while non-microtubule-invaded spines were more likely to be pruned after L-Dopa treatment (20 ± 3 % of spines, Fig. 4 F, G; white circles on the pruned spines).

In summary, our results indicate that abnormal L-Dopa-microtubule dynamics disrupts microtubule entry into spines, thereby reducing the ability of dendritic spines to resist pruning and consequently diminishing the number of the remaining dendritic spines. Altogether, the abnormal behavior of L-Dopa microtubules leads to a reduced ability of dendritic spines to establish mature synapses.

## DISCUSSION

This study describes for the first time the effects of L-Dopa, the standard treatment for Parkinson’s disease, on the density of dendritic spines and excitatory synapses of hippocampal cultured neurons and possible implications for synaptic dysfunction. We demonstrated that L-Dopa incorporation into microtubules reduced tyrosinated tubulin levels in favor of the appearance of a new post-translational modified form of microtubules composed of L-Dopa-tubulin. This alteration disrupted normal microtubule dynamics in the dendritic shaft, critical for maintaining dendritic spine density, and led to a reduction in the mature forms of dendritic spines in wild type hippocampal neurons. In these neurons, L-Dopa treatment not only reduced dendritic spine density, but also affected the percentage of excitatory synapses in the remaining spines, which could potentially affect synaptic plasticity and neuronal network function.

The deleterious effect of L-Dopa treatment could be due to the alteration of many cellular mechanisms, but the use of the knockout models for key enzymes involved in the tubulin detyrosination/tyrosination cycle confirmed that the observed effects were indeed mediated by L-Dopa incorporation into microtubules. We showed that in neurons from mice models in which L-Dopa incorporation is impaired, due to the absence of the ligase enzyme (TTL KO) or the reduction in the deTyrosinated tubulin pool that acts as substrate (SVBP KO), dendritic spine and synapse density was unaffected even in the presence of L-Dopa.

L-Dopa treatment altered global microtubule dynamics only in wild type neurons, increasing catastrophe frequency and reducing comet lifetime. These changes likely contributed to the observed reduction in microtubule invasion into dendritic spines, increasing spine vulnerability. Spines not invaded by microtubules were more prone to pruning, confirming the protective role of microtubule invasion in spine stability. These observations were similar to those described in other pathological contexts, such as Alzheimer’s disease neuronal model, in which invaded spines were more resistant to Aβ-induced synaptotoxicity (Peris, Parato, Qu, Soleilhac, Lante, et al., 2022).

Parkinson’s disease results from neuronal loss in the substantia nigra, and it is considered the second most common neurodegenerative disease, affecting 2-3% of the human population over the age of 65. Among antiparkinsonian therapies, the administration of L-Dopa shows the greatest symptomatic efficacy. Over time, almost all Parkinson’s patients require treatment with this tyrosine analog (LeWitt & Fahn, 2016; PD MED Collaborative Group, 2014). However, chronic long-term treatment can lead to various motor and nonmotor complications such as levodopa-induced dyskinesia and neuropsychiatric complications such as dopamine dysregulation syndrome (DDS) (Aquino & Fox, 2015; Beaulieu-Boire & Lang, 2015). L-Dopa-induced-dyskinesia has been associated with different events including pulsatile stimulation of dopamine (DA) receptors and abnormalities in non-dopaminergic transmitter systems. These elements together lead to alterations in the neuronal firing patterns that communicate between the basal ganglia and the cortex (Bastide et al., 2015). Additionally, alterations in dendritic spine density in medium spiny neurons have also been observed (Gomez et al., 2019).

In previous work, we described that the incorporation of L-Dopa into tubulin induces several changes that could compromise neuronal integrity. For example, we observed that the trafficking and distribution of mitochondria along the axons is affected, which could lead to a reduction in ATP availability, affecting the energy requirements of the neuron (Zorgniotti et al., 2021). In the present work, we also found that the incorporation of L-Dopa into tubulin affects the number of dendritic spines, their maintenance and the formation and consolidation of mature synapses.

We have previously described, using protein cellular extracts and purified tubulin in an in vitro system, that the incorporation of L-Dopa into tubulin was irreversible (Zorgniotti et al., 2021). We have now confirmed that the VASH1-SVBP complex is less efficient at releasing L-Dopa than tyrosine from microtubules due to alteration in the binding parameters of the enzyme induced by the presence of L-Dopa on the microtubular lattice. This observation allows us to speculate that, after years of administration, L-Dopa could potentially accumulate on α-tubulin and progressively replace dynamic tyrosinated microtubules in all neuronal cells, leading to the progressive chronic accumulation of L-Dopa-tubulin on synaptic microtubules, altering synaptic transmission and accelerating neurodegeneration.

It is still unknown if the presence of L-Dopa-microtubules or the reduction of Tyr-microtubules is the pathological feature that triggers synaptotoxicity. It has been shown that a balanced detyrosination/tyrosination tubulin cycle is necessary for the maintenance of synaptic plasticity. Moreover, increased microtubule tyrosination restored microtubule entry into spines suppressing the loss of synapses induced by amyloid-β peptide (Peris, Parato, Qu, Soleilhac, Lante, et al., 2022). Thus, the presence of L-Dopa-tubulin would unbalance the fragile equilibrium between tyrosinated and detyrosinated microtubules essential in the synaptic compartment, inducing synaptic dysfunction and damage.

Extensive studies will be necessary to elucidate this question, and to assess the impact of L-Dopa incorporation into microtubules on the synaptic compartment in vivo, as well as its potential role in L-Dopa-induced dyskinesia. Nevertheless, our results could lay the foundation for a new perspective on the pathogenesis of levodopa-induced dyskinesia in patients with Parkinson’s disease, identifying tubulin detyrosination/tyrosination as a key player in the regulation of synaptic transmission and neuronal function, and introducing the detyrosination/tyrosination cycle enzymes and microtubule dynamics as potential targets for therapeutic intervention.

## ACKNOWLEDGEMENTS.

We thank E Borel, K Vargas, S Ismail, Y Goldberg and A Buisson for helping with neuronal cultures and scientific discussion, C Ramel and people from the animal facilities for animal care; P Meresse for help in lentivirus preparation, C Bosc and B Blot for helping with molecular biology, Y Saoudi from PIC-GIN, for helping us with confocal microscopy. We are grateful to M Decressac for engaging in valuable scientific discussions that greatly contributed to this work. This work was performed at the Photonic Imaging Center of Grenoble Institute Neuroscience (PIC-GIN, Univ Grenoble Alpes – Inserm U1216) which is part of the ISdV core facility and certified by the IBiSA label.

## FUNDING

This work was supported by the doctoral fellowship from the Consejo Nacional de Investigaciones Científicas y Técnicas (CONICET-Argentina) and by France Parkinson post-doctoral fellowship (2023-CS-0005) to A. Zorgniotti; France Alzheimer (SynapCyAlz AAP PFA 2022 R23052CC) and Fondation Recherche Alzheimer (C7H-PRI23A64 - FRAlz) to L. Peris; Agence National de la Recherche (SPEED-Y ANR-20-CE16-0021), the Leducq Foundation (research grant no. 20CVD01) and NeuroCAP-Servier grants to M.-J.Moutin; The Company of Biologist and the Journal of Cell Science travel grant to A. Zorgniotti. Y. Ditamo and C.G.Bisig, were supported by the Agencia Nacional de Promoción Científica y Tecnológica de la Secretaria de Ciencia y Tecnología del Ministerio de Cultura y Educación (Prestamo BID-PICT 2019-1584), the Consejo Nacional de Investigaciones Científicas y Técnicas (CONICET-Argentina), and the Secretaria de Ciencia y Técnica from the National University of Córdoba-Argentina.

## CONTRIBUTIONS

AZ, GB and LP conceived and designed the study.

AZ, AS and MA performed molecular biology experiments and lentivirus preparations.

MJM and LP supervised mice production.

AZ, AS and LP performed experiments to analyze spine density and excitatory synapses in mice hippocampal cultured neurons.

AZ and AS performed biochemical experiments with neuronal samples.

AZ, CS, SRR and MJM conceived and performed in vitro TIRF and immunofluorescence studies.

AZ, GB and LP performed the associated statistical analysis.

AZ and LP designed and performed analysis of microtubule entry into dendritic spines and dendritic MT dynamics.

AZ, YD, GB and LP wrote the manuscript, with contributions from all co-authors.

## Declaration of Competing interests

The authors declare no competing interests.

## Abbreviations

MT: microtubule
TTL: tubulin tyrosine ligase
VASH-SVBP: vasohihin-small vasohibin binding protein
PD: Parkinson’s disease
LID: L-Dopa-induced-dyskinesia

## Notes

### Competing Interest Statement

The authors have declared no competing interest.

