## Supplemental Figures for "L-Dopa incorporation into tubulin alters microtubule dynamics and reduces dendritic spine invasion and synapse maintenance"

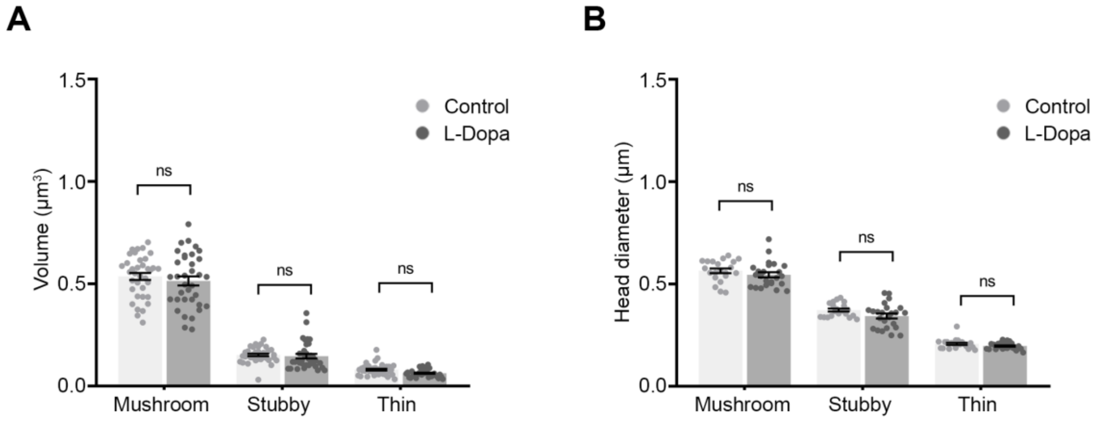


**Supplementary Figure 1 L-Dopa induced dendritic spine density reduction does not modify spine morphology.** Morphological analysis of dendritic spines in eGFP expressing wild type (wild type) hippocampal neurons (18 DIV) treated either with 0.4 mM L-Dopa or the vehicle (Control). Analysis of dendritic spine head volume **(A)** and diameter **(B)**. Data represent mean ± SEM; n = 36, n = 35 control and L-Dopa-treated neurons, respectively from at least three different embryos and neuronal cultures. Mann-Whitney test, ns = not significant.

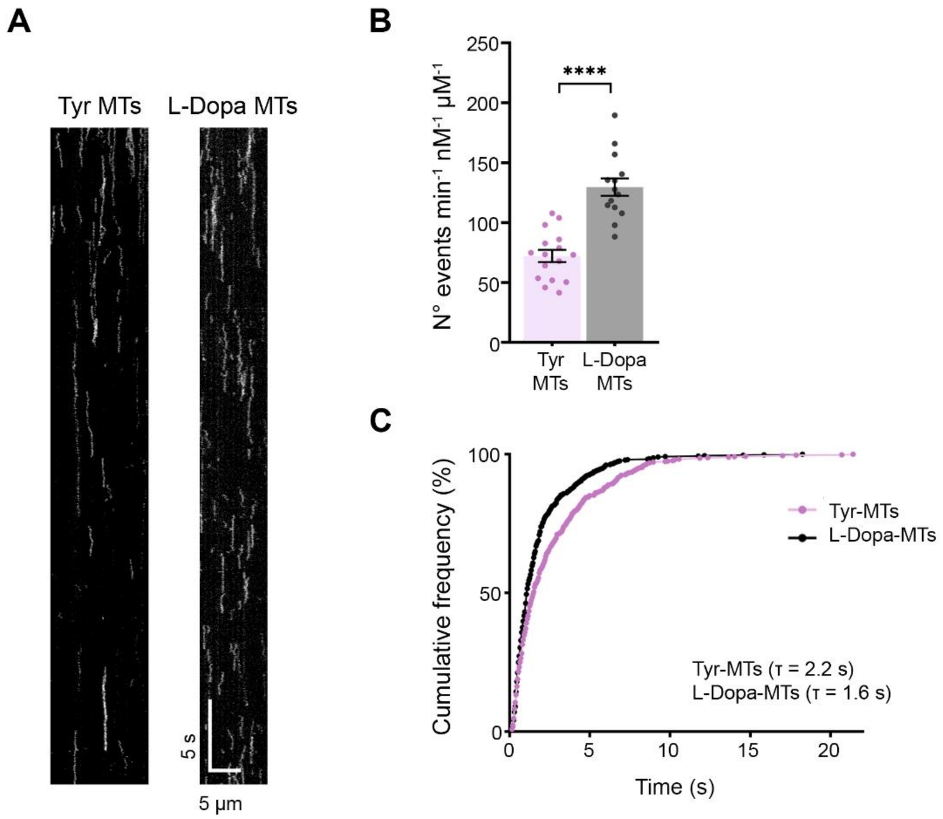


**Supplementary Figure 2 L-Dopa incorporation into microtubules perturbs VASH1-SVBP binding. (A)** Representative kymographs of single molecules of catalytically inactive sfGFP-tagged VASH1–SVBP at a concentration of 50 pM bound to Taxol-stabilized microtubules enriched in tyrosinated (Tyr MTs) or L-Dopa (L-Dopa MTs) tubulin. Scale bars: horizontal, 5 µm; vertical, 5 s. **(B)** Analysis of binding frequency. Number of binding events of the deadVASH1-SVBP enzyme complex to tyrosinated (Tyr MTs) or L-Dopa (L-Dopa MTs) microtubules. Each point represents an individual microtubule. Data represent mean ± SEM; n = 16 Tyr-MTs and n = 14 L-Dopa-MTs. Unpaired t-test; ****$P < 0.0001$. **(C)** Cumulative frequency of the residence times measured in TIRF movies taken during the 30 min following addition of enzyme complexes to tyrosinated (Tyr MTs) or L-Dopa (L-Dopa MTs) microtubules. The mean residence time (τ) is obtained by fitting the curve with a mono-exponential function.
